# STAT3 expression in dendritic cells protects mice from colitis by a gut microbiome-dependent mechanism

**DOI:** 10.1101/2021.07.31.453520

**Authors:** Jianyun Liu, Keely L. Szilágyi, Maegan L. Capitano, Abhirami K. Iyer, Jiefeng He, Matthew R. Olson, Jianguang Du, William Van Der Pol, Casey Morrow, Baohua Zhou, Mark H. Kaplan, Alexander L. Dent, Randy R. Brutkiewicz

## Abstract

An imbalance in gut homeostasis results in local and systemic pathogenesis. It is still not well-understood how the immune system interacts with the gut microbiome and maintains a delicate balance. Here, we utilized a mouse model in which STAT3 expression is deleted in CD11c^+^ (i.e., dendritic) cells (STAT3 cKO); these mice developed an ulcerative colitis-like disease, colon carcinoma and myelodysplastic syndrome-like disease. Circulating IgE levels in STAT3 cKO mice were significantly elevated. The gut microbiome was indispensable for the observed pathogenesis, as treatment with broad-spectrum antibiotics or cross-fostering STAT3 cKO pups with mothers harboring a different microbiome prevented disease development. Gut microbiome analyses suggested that decreased commensal bacteria and increased pathogenic bacteria most likely contributed to disease. Our data suggest that STAT3 controls the manifestation of inflammation in the gut caused by the microbiome. Therefore, we conclude that a deficiency of STAT3 in DCs is sufficient to trigger uncontrolled inflammation and the development of inflammatory bowel disease.

## Introduction

Intestinal homeostasis is a finely-tuned dynamic interplay between the gut microbiome and host immunity (1). An imbalance of this homeostasis results in inflammatory bowel disease (IBD), including Crohn’s disease and ulcerative colitis, as well as cancer (2). About 3.1 million adult Americans (i.e., 1.3% of the population) are estimated to have IBD (3). It is generally accepted that IBD is caused by improper immune responses against the gut microbiome due to unknown mechanisms. Current treatments for IBD can only mitigate the symptoms, rather than providing a cure (4). Intestinal dendritic cells (DCs) are important for maintaining intestinal homeostasis by inducing protective immunity against infectious agents and tolerance to harmless food antigens and the gut microbiome (5–7), but the mechanism is not well-understood. Further understanding of the interplay between the gut microbiome and intestinal DCs will provide a better therapeutic strategy for the treatment of IBD.

The Janus kinase–signal transducer and activator of transcription (JAK/STAT) signaling pathway provides a rapid response to receptor-ligand coupling and is essential for many biological processes (8, 9). STAT3 is a member of the STAT protein family and is phosphorylated by JAKs responding to cytokines and growth factors, such as IL-6, IL-10, LIF and hepatocyte growth factor (9, 10). Mutations in STAT3 cause autosomal dominant hyper-IgE syndrome (AD-HIES), a primary immunodeficiency with many immunological and non-immunological manifestations (11–13). *STAT3* mutations and polymorphisms have also been reported to link to IBD (14, 15). Research using mouse models suggests that abnormal STAT3 expression in immune cells results in IBD (16). However, the role of STAT3 in different mouse models can differ completely and is believed to be context-dependent. Thus, it is important to investigate how STAT3 contributes to IBD under different conditions.

In this study, we generated a transgenic mouse model specifically deleting STAT3 in CD11c+ cells (i.e., dendritic cells; DCs) and found that these mice spontaneously develop colitis and a high incidence of colon carcinoma. We further analyzed immunological responses in the diseased mice and how the gut microbiome contributes to disease development. This model could be very useful for the identification and study of effective IBD treatments.

## Results

### Deletion of STAT3 in DCs results in colitis, colon adenocarcinoma, elevated IgE and a myelodysplastic syndrome (MDS)-like disease

We have previously reported that STAT3 signaling promotes CD1d-mediated lipid antigen presentation to natural killer T (NKT) cells (17). To study how a DC-specific deficiency in STAT3 impacts NKT cell function *in vivo*, we generated *Stat3* conditional KO (STAT3 cKO) mice by back-crossing *Stat3*^fl/fl^ mice to CD11c-Cre (*Itgax*-Cre) mice. A deficiency in DC STAT3 from STAT3 cKO mice was confirmed by qPCR, flow cytometry and Western blot analyses (Fig. 1A-1C, Fig. S1A-C). By comparison, STAT3 expression in other cell types remained intact (Fig. S1D-E). When examining these mice, we serendipitously found that ∼70% of STAT3 cKO mice died by 30 weeks of age (median survival time=18.5 weeks, Fig. 1D) due to humane endpoints being reached from rectal prolapse or sudden death. These STAT3 cKO mice, but not their co-housed WT/het littermates, developed lymphocytic colitis with enlarged mesenteric lymph nodes (Fig. 1E-F). About two thirds of these mice (4 out of 6 mice examined) also developed colon adenocarcinoma (Fig. 1G). Furthermore, the histologic scores of large intestines from STAT3 cKO mice were significantly higher than those from WT mice (Fig. 1H). Histology of the small intestines from STAT3 cKO mice appeared to be normal (data not shown).

**Figure 1.**
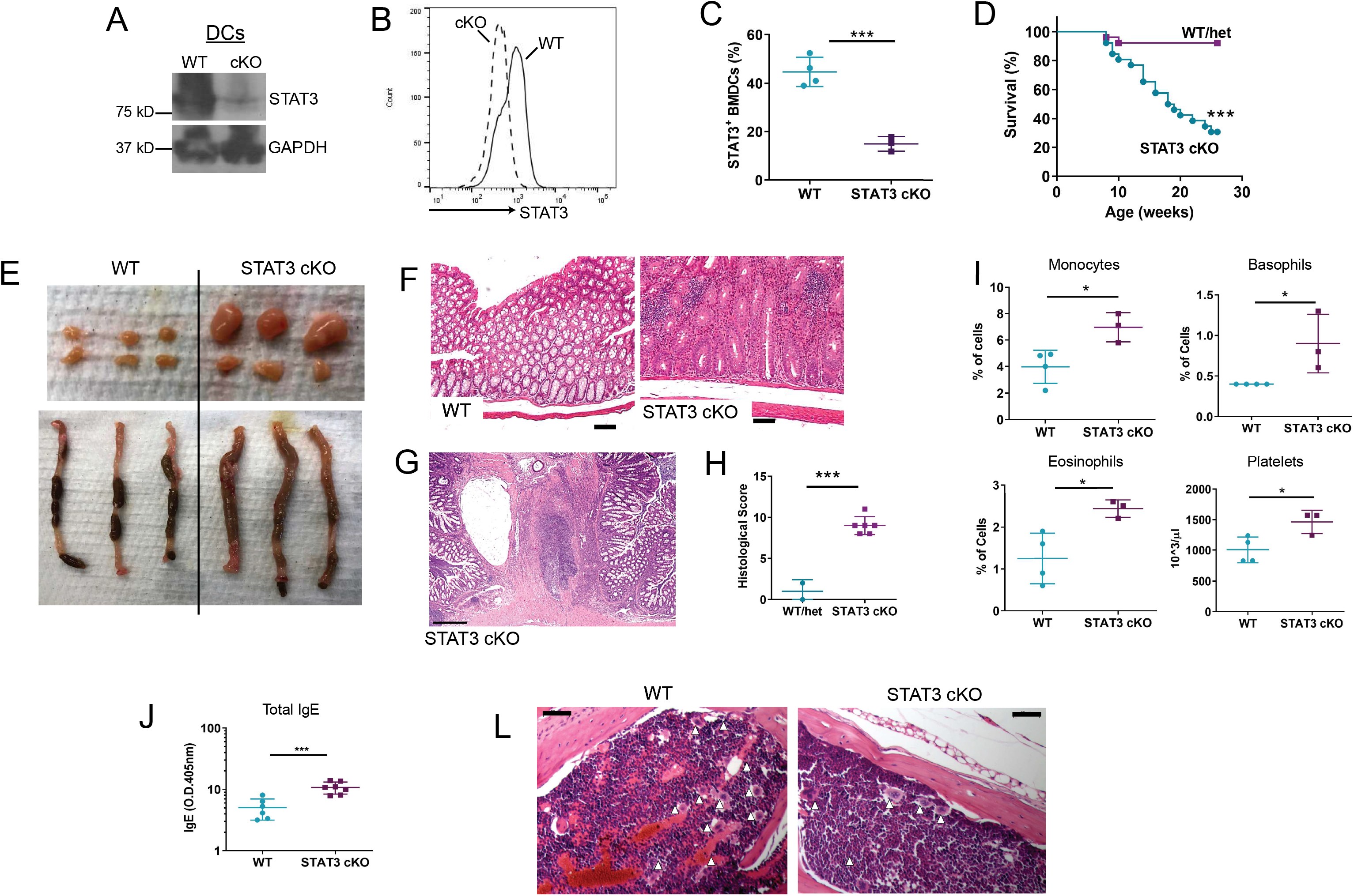
STAT3 cKO mice develop colitis, colon adenocarcinoma, elevated IgE and an MDS-like disease. The expression of STAT3 in bone marrow-derived dendritic cells (BMDCs) from one representative STAT3 cKO mouse and a WT littermate was analyzed by Western Blot (A) or flow cytometry (B). Percentages of STAT3-positive BMDCs from multiple animals are summarized in (C). (D) Survival rates of STAT3 cKO mice and their WT/het littermates. (E) Mesenteric lymph nodes (top) and colons (bottom). (F) H&E staining of cecum/colon tissues from STAT3 cKO mice and their WT littermates. The black bars denote 100 µm. (G) H&E staining of cecum/colon tissues from STAT3 cKO mice showing colon carcinoma. The black bar denotes 500 µm. (H) Summary of histology scores of cecum/colon tissues. (I) The levels of circulating monocytes, basophils, eosinophils, platelets, and (J) total IgE from STAT3 cKO mice and WT/het littermates. (K) H&E staining of sternal bone marrow from STAT3 cKO mice and their WT littermates. The black bars denote 50 µm. The white triangles point to megakaryocytes. The data are plotted as the mean ± SD. Each dot represents an individual animal. *, *p*<0.05; ***, *p*<0.001. Mann–Whitney unpaired *t* test (C, H, I, J) and log-rank test (D).

Blood cell profiling analysis demonstrated that STAT3 cKO mice had higher levels of circulating monocytes and granulocytes, as well as platelets (Fig. 1I). Total IgE in the sera of STAT3 cKO mice was also elevated compared to their WT/het littermates (Fig. 1J). Histological analysis of the bone marrow suggested that STAT3 cKO mice developed a myelodysplastic syndrome (MDS)-like disease based on the presence of dysplastic megakaryocytes (Fig. 1K). Thus, a deficiency of STAT3 in DCs resulted in the development of an ulcerative colitis-like disease, colon cancer and MDS-like disease in STAT3 cKO mice.

### STAT3 cKO mice have increased Th17 cells

With the substantial inflammation present in STAT3 cKO mice, it was important to assess the different pro-inflammatory cytokines that could have contributed to this disease state. Thus, we examined circulating cytokines in the sera of STAT3 cKO mice with a multiplex bead array. Among the 32 cytokines/chemokines we examined, we found that STAT3 cKO mice had elevated levels of circulating IFN-γ, IL-10, IL-17, IP-10, TNF-α and MIP 3α (Fig. 2A and 2B). STAT3 cKO mice had fewer splenic T, B and NK cells, but more DCs and macrophages (Fig. S2A-C). Interestingly, STAT3 cKO mice had more Th17 cells in the spleen, liver, MLN and colon (Fig. 2C), with the exception of the Peyer’s Patch (Fig. S2F). Neither Th1 cells (Fig. S2D-E) nor Tregs (Fig. S2G-H) were found to be higher in STAT3 cKO mice. Therefore, our data demonstrate that STAT3 cKO mice exhibit a Th17-biased colitis.

**Figure 2.**
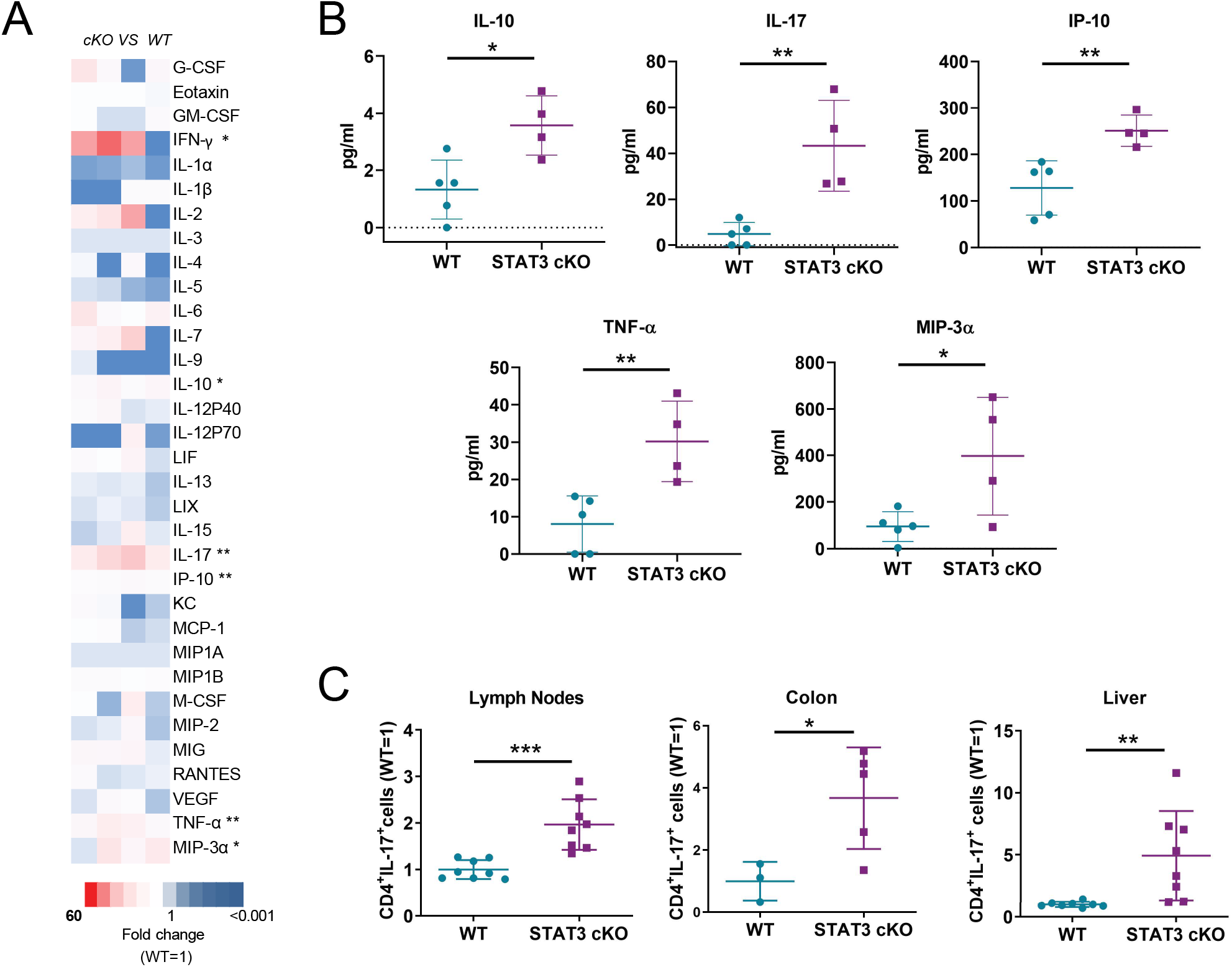
STAT3 cKO mice develop a Th17-biased colitis. A) A heat-map showing the relative levels of circulating cytokines and chemokines in STAT3 cKO mice compared to WT littermates (WT=1). The expression levels of cytokines and chemokines that are significantly increased in STAT3 cKO mice are shown in (B). (C) The levels of Th17 cells (CD3^+^CD4^+^IL-17A^+^) from mesenteric LNs, colons and livers relative to WT are summarized. The data are plotted as the mean ± SD. Each symbol represents an individual animal. *, *p*<0.05; **, *p*<0.01; ***, *p*<0.001. Mann–Whitney unpaired *t* test.

### Antibiotic treatment alleviates gastrointestinal disease in STAT3 cKO mice

Studies have shown that the microbiome likely contributes to the development of colitis (18). To assess the possibility that the microbiome contributed to the observed colitis, STAT3 cKO mice and their WT/het littermates were given medicated chow containing broad-spectrum antibiotics (Uniprim) for 10 weeks. Such treated STAT3 cKO mice exhibited no disease symptoms (Fig. 3A-B). STAT3 cKO mice on regular feed showed significantly higher levels of neutrophils and lower numbers of lymphocytes in their blood compared to their WT/het littermates. In contrast, STAT3 cKO mice on Uniprim feed for 10 weeks had equivalent levels of neutrophils and lymphocytes as their WT/het littermates (Fig. 3C-D), and normal levels of circulating IgE (Fig. 3E). Notably, however, even after 10 weeks of antibiotic treatment, STAT3 cKO mice still had more Th17 cells present in the gut as compared to their WT/het littermates (Fig. 3F), suggesting this cell population is not a major contributor to colitis under these conditions. Nonetheless, we conclude that the development of colitis in STAT3 cKO mice is dependent on the gut microbiome.

**Figure 3.**
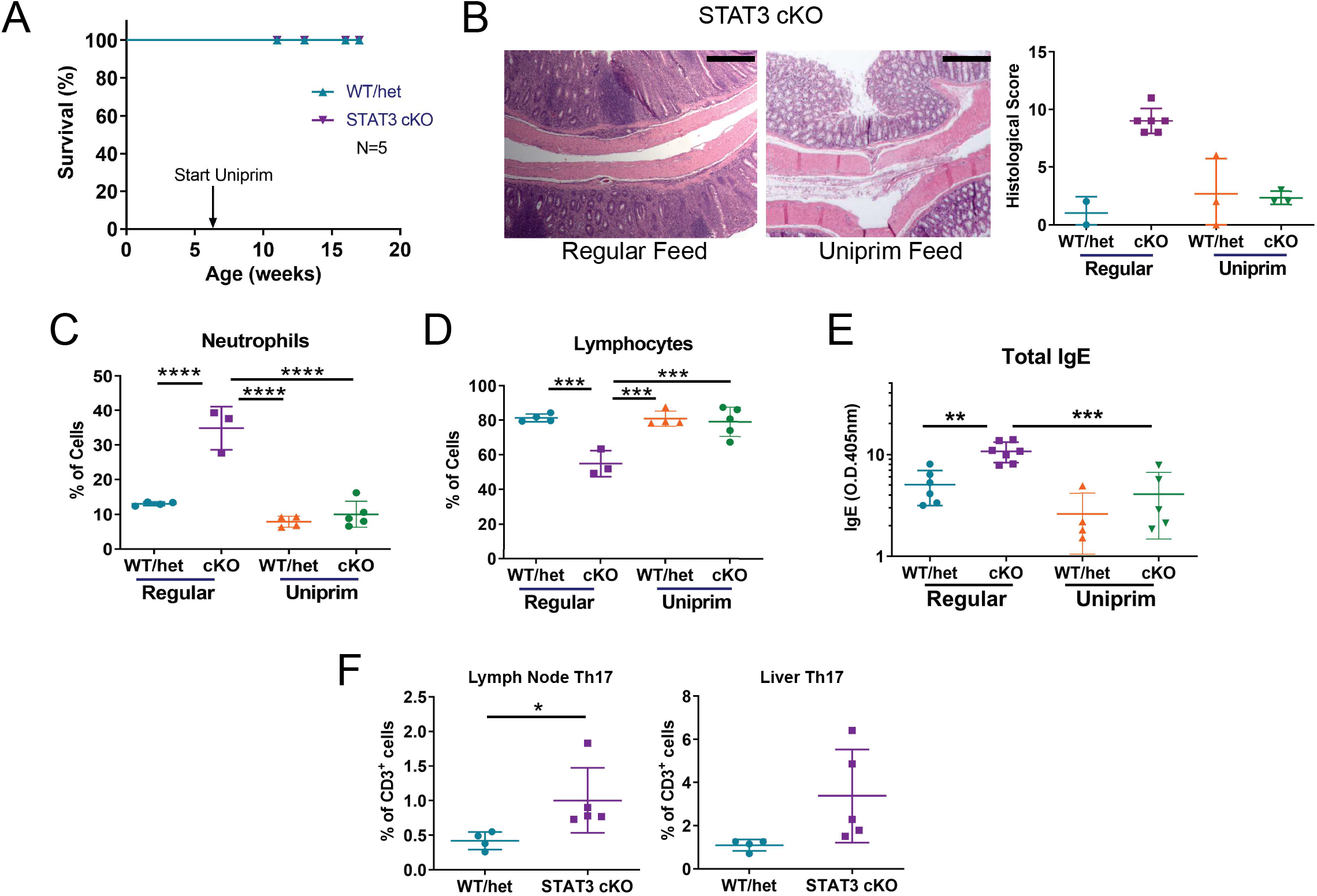
Antibiotic treatment alleviates disease in STAT3 cKO mice. (A) STAT3 cKO mice and their WT/het littermates were given Uniprim-medicated feed *ad libitum* for 10 weeks and their survival rates were determined. (B) H&E staining of the cecum/colon from STAT3 cKO mice on normal or Uniprim feed. The black bars denote 500 µm. The histology scores from STAT3 cKO mice and their WT/littermates on regular or Uniprim feed are summarized. Each symbol represents an individual animal. The levels of neutrophils (C), lymphocytes (D) and total IgE (E) from STAT3 cKO and WT/het littermates on regular or Uniprim feed are shown. (F) STAT3 cKO mice and WT/het littermate controls were kept on Uniprim feed for 10 weeks. IL-17-producing CD4^+^ T cells in mesenteric LNs and liver MNCs were analyzed and are shown. The data are plotted as the mean ± SD. Each symbol represents an individual animal. *, *p*<0.05; **, *p*<0.01; ***, *p*<0.001. Log-rank test (A), one-way ANOVA test (C-E) and Mann–Whitney unpaired t test (F).

### The MDS-like disease in STAT3 cKO mice is reversible

Besides colitis and colon cancer, STAT3 cKO mice also exhibited an MDS-like disease (Fig. 1K). Bone marrow from mice fed on regular chow looked pale (Fig. 4A). Further analysis by flow cytometry showed that there were more lineage^−^ Sca-1^+^cKit^+^ (LSK) cells and fewer megakaryocyte/erythrocyte progenitors (MEP, Lin^−^ Sca-1^−^ c-Kit^+^ FcγII/IIIR^−^ CD34^+/-^) in STAT3 cKO mice compared to their WT/het littermates (Fig. 4C), providing a possible explanation for the fewer megakaryocytes observed in the bone marrow of STAT3 cKO mice (Fig. 1K). Although not significantly different, there was a trend of increases in granulocyte/monocyte progenitor cells (GMP, Lin^−^ Sca-1^−^ c-Kit^+^ FcγII/IIIR^hi^ CD34^+/-^) (Fig. 4C), suggesting a skewing of myelopoiesis toward the production of GMPs over MEPs. However, to our surprise, after being on the Uniprim diet for 6-10 weeks, bone marrow cells in STAT3 cKO mice became normal and showed no differences from their WT/het littermates (Fig. 4D & E). Therefore, we conclude that the MDS-like disease in STAT3 cKO mice is reversible.

**Figure 4.**
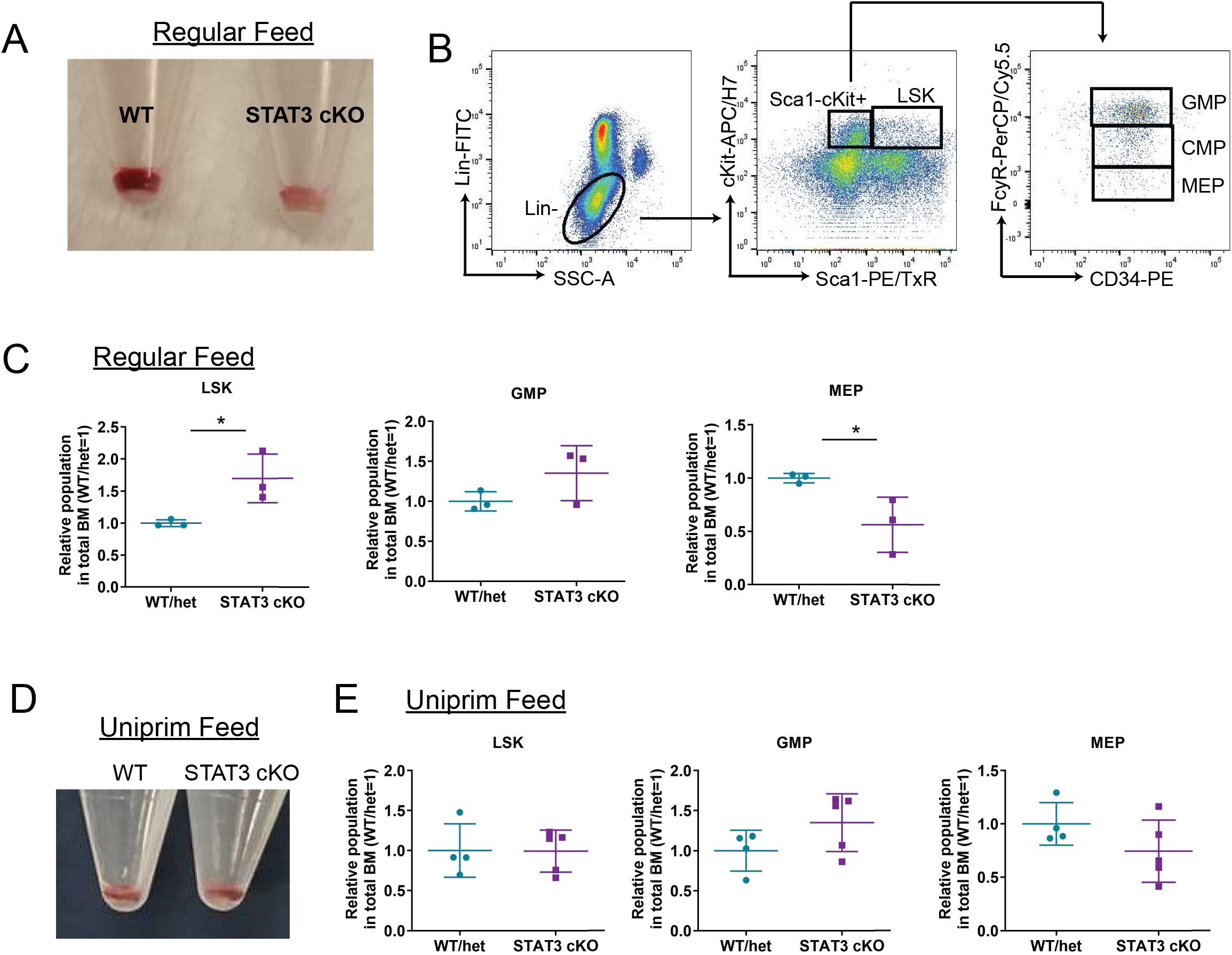
Normal bone marrow cells in Uniprim-treated STAT3 cKO mice. (A) Bone marrow was harvested from STAT3 cKO mice and their WT littermates by flushing the femur and tibia. (B) Bone marrow cells from STAT3 cKO mice were stained with the indicated antibodies and the gating strategy for hematopoietic stem cells is shown. (C) Bone marrow cells from mice fed with normal chow were stained with antibodies specific for the indicated hematopoietic stem cell subpopulations. The relative levels of LSK, GMP and MEP in total bone marrow were normalized and graphed (WT/het=1). (D and E) Bone marrow cells from mice that were fed Uniprim chow were harvested and stained with antibodies specific for the indicated hematopoietic stem cell subpopulations. The relative levels of LSK, GMP and MEP in the total bone marrow were normalized and graphed (WT/het=1). The data are plotted as the mean ± SD. Each dot represents an individual animal. *, *p*<0.05. Mann–Whitney unpaired *t* test.

### Altered gut microbiome in diseased STAT3 cKO mice

To further address the possibility that the colitis in STAT3 cKO mice was gut microbiome-dependent, we constituted STAT3 cKO mice and their co-housed WT/het littermates with a different gut microbiome (GM) by cross-fostering neonatal mice to wildtype mothers obtained from The Jackson Laboratory (JAX). To directly test whether there were differences in the gut microbiome between the groups of mice not being cross-fostered (GM-IUSM) versus those that were (GM-JAX), fecal pellets from each group of mice were analyzed by 16S rRNA sequencing microbiome analysis. The overall compositional difference of the microbiome across all samples at the family level is shown in the heat-map in Fig. 5A. The beta diversities of each mouse sample analyzed were visualized on Principal Coordinate Analysis (PCoA) plots to indicate the similarities between samples (Fig. 5B). The data suggest that the co-housed GM-JAX WT and GM-JAX STAT3 cKO littermates harbored a similar microbiome, as expected. The microbiome in individual GM-IUSM WT mice, although different from that in GM-JAX mice, was similar within the group. However, the microbiome in GM-IUSM STAT3 cKO mice diverged from those of their co-housed WT littermates. The gut microbiome in individual GM-IUSM STAT3 cKO mice was also distinct from each other and scattered on the PCoA plots (Fig. 5B), suggesting there was substantial variation in the intestinal microbiome community among these diseased mice.

**Figure 5.**
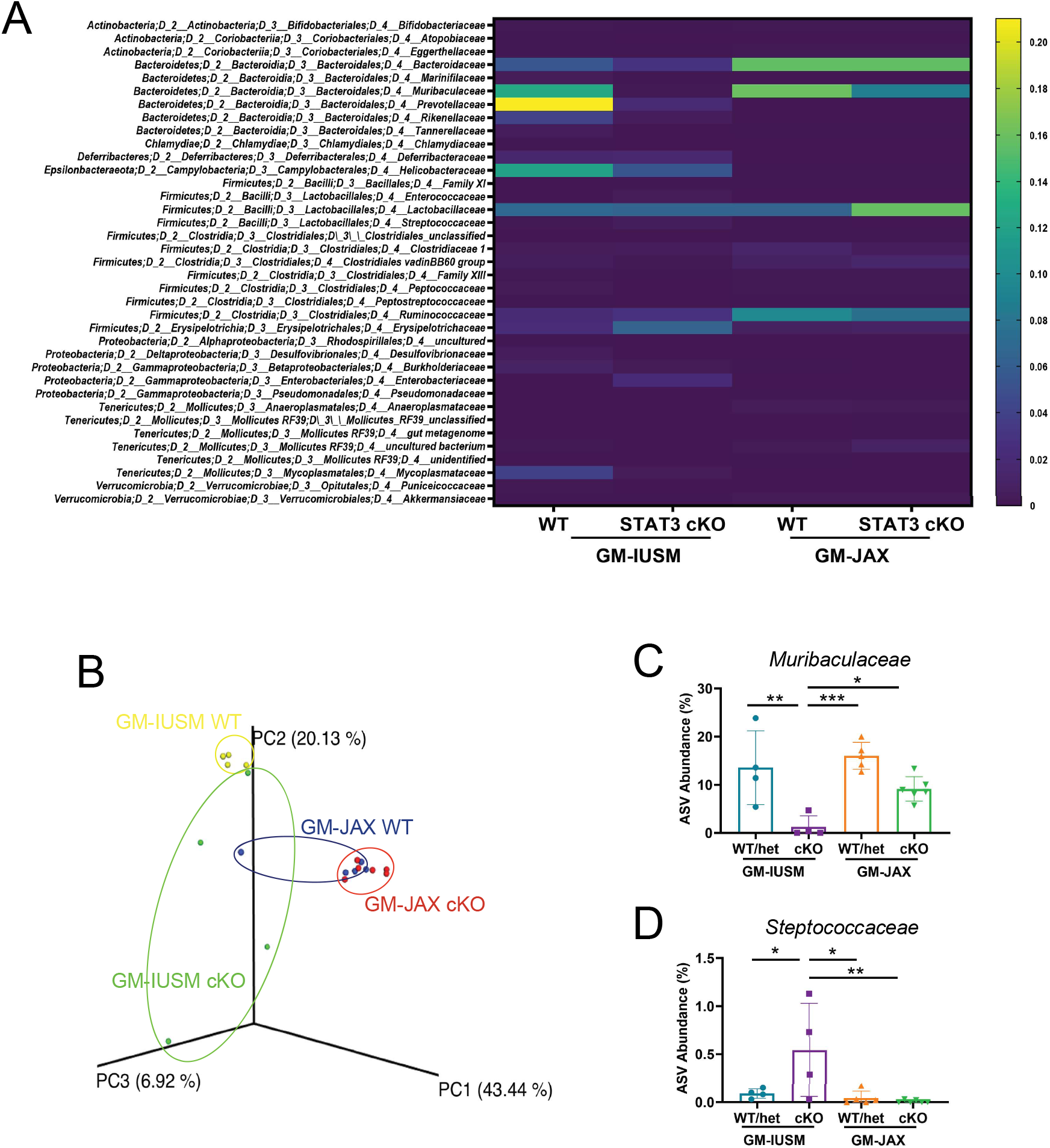
Altered gut microbiome in GM-IUSM STAT3 cKO mice. (A) Fecal pellets from GM-IUSM and GM-JAX STAT3 cKO mice and their WT/het littermates were analyzed by 16S rRNA sequencing analysis. Amplicon Sequence Variants (ASV) at the family level were graphed as a heatmap. (B) PCoA of the variation of microbial communities between samples. The ASV values of *Muribaculaceae* (C) and *Steptococcaceae* (D) are shown. The data are plotted as the mean ± SD. Each symbol represents an individual animal. *, *p*<0.05; **, *p*<0.01; ***, *p*<0.001. One-way ANOVA test.

We closely examined the difference in bacterial composition in the gut microbiomes of STAT3 cKO mice and their co-housed WT/het littermates. *Pseudomonadaceae* and *Anaeroplasmataceae* were found in the microbiomes of GM-JAX mice, but not in GM-IUSM mice (Fig. S3A). By contrast, the microbiome from GM-IUSM mice had more *Deferribacteraceae*, *Helicobacteraceae* and *Mycoplasmataceae* (Fig. S3B). *Rikenellaceae* and *Prevotellaceae*, although absent in GM-JAX mice, were more abundant in GM-IUSM WT mice than GM-IUSM STAT3 cKO mice (Fig. S3C). *Muribaculaceae*, a common and abundant inhabitant in the normal gut flora of mice (19), was significantly reduced in GM-IUSM STAT3 cKO mice (Fig. 5C). Furthermore, GM-IUSM STAT3 cKO mice had more potential pathogenic bacteria of the *Streptococcaceae*, *Enterococcaceae* and *Enterobacteriaceae* families (Fig. 5D, Fig. S3D). Overall, the gut microbiome in GM-IUSM STAT3 cKO mice was distinctly different from their WT/het littermates and potentially associated with a decreased abundance of commensal bacteria and increased pathogenic bacteria.

### GM-JAX STAT3 cKO mice do not develop colitis

As shown by the microbiome analysis, GM-JAX STAT3 cKO mice had a similar gut microbiome as their WT/het littermates (Fig. 5A and 5B). Additionally, the GM-JAX STAT3 cKO mice also did not exhibit any disease symptoms (Fig. 6A). indicating that the differences in the microbiome contributed to disease in STAT3 cKO mice with the GM-IUSM. GM-JAX STAT3 cKO mice also had normal levels of circulating total IgE and IL-17 (Fig. 6B), as well as Th17 cells (Fig. 6C). Moreover, whereas GM-IUSM STAT3 cKO were not able to breed due to severe disease development, GM-JAX STAT3 cKO mice could breed normally. Additionally, the histology of bone marrow from GM-JAX STAT3 cKO mice was also no different from their WT/het littermates (Fig. 6D). Therefore, our results demonstrate that the development of an ulcerative colitis-like disease in STAT3 cKO mice is dependent on the gut microbiome.

**Figure 6.**
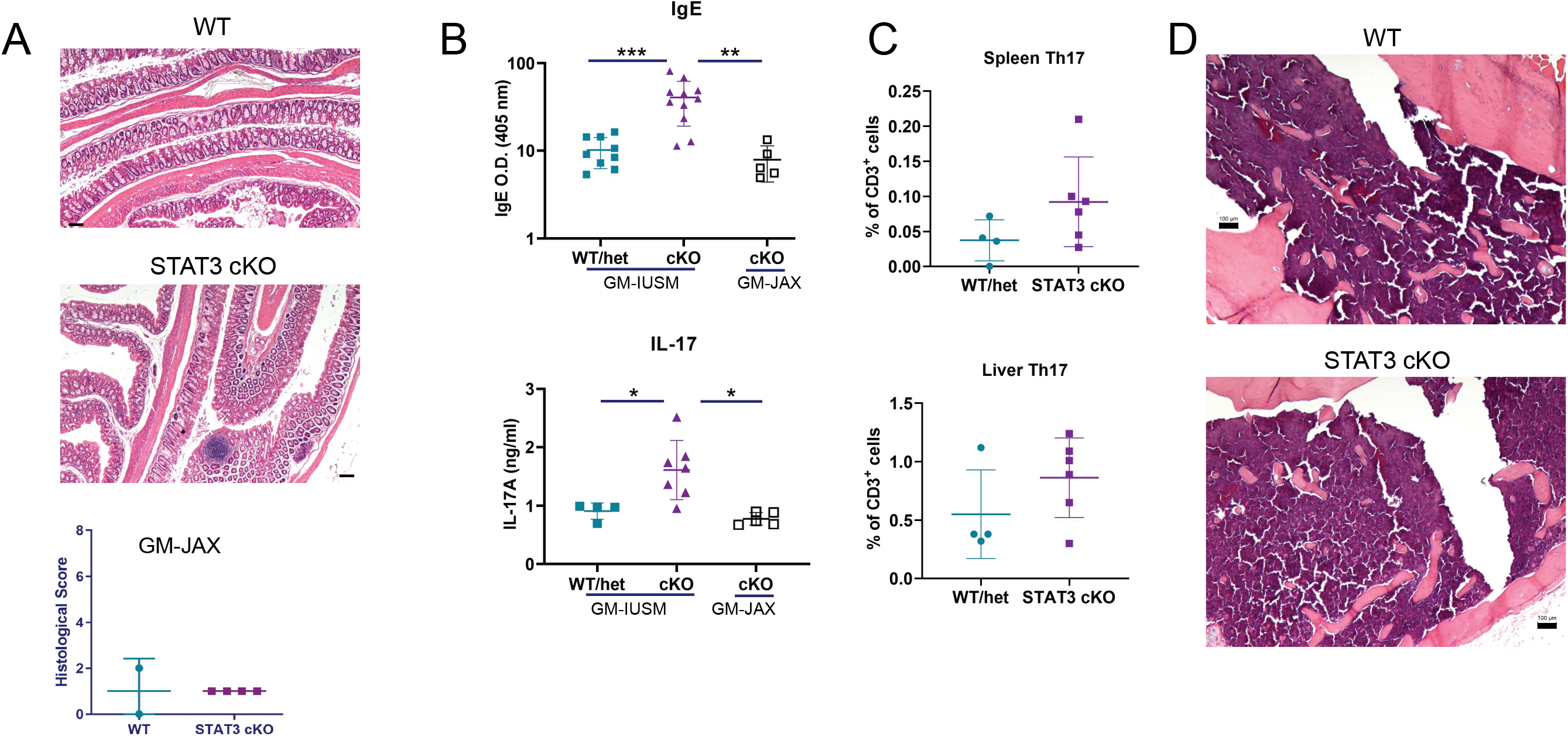
GM-JAX STAT3 cKO mice do not develop colitis. (A) H&E staining of cecum/colon tissues from STAT3 cKO mice and WT/het littermates with the JAX gut microbiome (GM-JAX). The histology scores are summarized below the images. The black bars denote 100 µm. (B) Serum IgE and IL-17 from GM-JAX STAT3 cKO mice were measured and compared to GM-IUSM WT/het and STAT3 cKO mice. (C) Quantitation of spleen and liver Th17 cells from GM-JAX STAT3 cKO mice and their WT/het littermates. (D) H&E staining of femoral bone marrow from GM-JAX STAT3 cKO mice and their WT littermates. The black bars denote 100 µm. The data are plotted as the mean ± SD. Each symbol represents an individual animal. *, *p*<0.05; **, *p*<0.01; ***, *p*<0.001. One-way ANOVA test.

**Figure 7.** Graphical illustration showing that both a deletion of STAT3 in DCs and the presence of certain bacteria in the gut microbiome promote the development of colitis and colon cancer in mice.

## Discussion

In the current study, we show direct evidence that a deficiency in STAT3 signaling in DCs results in Th17-biased gut inflammation, colitis and colon cancer via a gut microbiome-dependent mechanism. We only found inflammation and pathology in the large intestine (cecum and colon) in GM-IUSM STAT3 cKO mice; the small intestines from these mice appeared to be normal and no increased levels of Th17 cells were observed in their Peyer’s patches. In a prior study, a deficiency of STAT3 in Treg cells was shown to result in spontaneous Th17-biased colitis, which could be lessened by treatment with neutralizing anti-IL-17 antibody (20). We observed comparable numbers of Tregs in GM-IUSM STAT3 cKO and WT/het mice; this suggests that colitis in the STAT3 cKO mice was not caused by impaired Treg development or priming. Normal Treg differentiation and function, but less tolerogenic DCs were also previously reported in AD-HIES patients (21).

IL-10 is an important immunomodulator for maintaining intestinal homeostasis. Similar to previous reports (22–24), we also observed colitis in IL-10 KO mice. Considering the elevated level of IL-10 in STAT3 cKO mice (Fig. 2B), this suggested that the colitis symptoms in these mice were not due to an IL-10 deficiency. As STAT3 is a downstream mediator of IL-10 signaling (25), our data indicate that impaired IL-10/STAT3 signaling in DCs causes colitis/gut inflammation. Moreover, IL-10 KO mice did not display elevated IL-17 or IgE (Fig. S4A-B) as STAT3 cKO mice do, suggesting that a lack of STAT3 in DCs was not equivalent to an IL-10 deficiency.

Our data suggest that the gut microbiome in GM-IUSM STAT3 cKO mice was linked to a decreased abundance of commensal bacteria and increased pathogenic bacteria. *Helicobacter spp*, a well-known gut bacterium that causes colitis in immunocompromised mice (26, 27), was detected in GM-IUSM mice, but not in mice purchased from The Jackson Laboratory. Notably, GM-JAX STAT3 cKO mice lacking *Helicobacter spp* in the gut microbiome, were in overall good health. However, in addition to *Helicobacter spp*, there are also other potential pathogenic bacteria, such as *Streptococcaceae*, *Enterococcaceae* and *Enterobacteriaceae*, which were also present in the microbiomes of GM-IUSM STAT3 cKO mice. Overrepresentation of *Enterococcaceae* and *Enterobacteriaceae* has been reported to be associated with IBD (28), hospitalizations (29) and Parkinson’s disease (30). Some bacteria, such as *Rikenellaceae*, *Prevotellaceae* and *Muribaculaceae*, appeared to be lacking in GM-IUSM STAT3 cKO mice. It is not clear whether these bacteria are protective against colitis, or if their absence in GM-IUSM STAT3 cKO mice is a result of inflammation. Previous work has shown that inflammation can alter the microbiome (22). *Rikenellaceae* has been reported to be reduced in the gut microbiomes of patients with non-alcoholic fatty liver disease (31, 32). Certainly, even in the presence of *Helicobacter spp*, different gut microbiomes can modulate disease phenotypes (33).

Interestingly, our PCoA analysis of the gut microbiome demonstrated that GM-JAX mice and GM-IUSM WT mice form tight clusters within the groups. However, the data from GM-IUSM STAT3 cKO mice were scattered, suggesting substantial variation in the intestinal microbiome community among these mice. It seems likely that there is a set of factors contributing to a healthy microbiome and the lack of any of these factors in GM-IUSM STAT3 cKO mice results in inflammation and colitis. The considerable variation in the gut microbiome of different co-housed mice may also be explained by gut inflammation in GM-IUSM STAT3 cKO mice as inflammation can alter the microbiome (22).

Bone marrow cells from Uniprim-fed STAT3 cKO mice were normal and were no different than their WT/het littermates (Fig. 4D and 4E), suggesting the MDS-like disease in STAT3 cKO mice is preventable. Interestingly, a vitamin B12 deficiency has been reported to cause MDS-like disease and the symptoms are reversible (34–36). A recent case report described a young man with gastrointestinal disease who presented with an MDS-like malady but experienced a rapid, back to normal bone marrow recovery after nutrient supplementation (37). The severe gut inflammation in STAT3 cKO mice could affect normal gut function, resulting in malnutrition in these mice. Therefore, the MDS-like syndrome in STAT3 cKO mice could have been caused by malnutrition and was thus reversible, once the gut inflammation was mitigated by Uniprim treatment.

A prior mouse model was generated to mimic the mutations in STAT3 found in AD-HIES patients (38). The mice carrying mutant STAT3 showed normal B cell development but elevated IgE. DCs from this mutant mouse model were also less responsive to IL-10. Naïve T cells from these STAT3 mutant mice could not differentiate into Th17 cells, due to dysfunctional STAT3, resembling the deficiency of Th17 cells in AD-HIES patients (39). AD-HIES patients are susceptible to lung and skin bacterial infections, but they do not usually develop IBD (40, 41). In a T cell-dependent mouse colitis model, STAT3 expression in T cells is required for the development of Th17-biased colitis (42). In the current study, we show that a STAT3-deficiency in DCs results in Th17-biased gut inflammation, colitis and colon cancer. Our data suggest that the colitis can develop when STAT3 is sufficient in T cells but deficient in DCs. Thus, it is likely that AD-HIES patients do not develop IBD because their T cells lack the the expression of functional STAT3. In addition, our observation that a STAT3 deficiency in DCs alone can elevate IgE levels in a gut microbiome-dependent manner, suggests that microbial stimuli also contribute to hyper-IgE in AD-HIES patients. Future work should focus on: 1) identifying the microbial stimuli in the gut microbiome that are causing inflammation in STAT3 cKO mice and 2) how DC STAT3 regulates gut homeostasis and IgE production *in vivo*.

In summary, we have generated a unique ulcerative colitis-like mouse model by selectively deleting STAT3 expression in DCs. These mice display different disease phenotypes depending upon the overall gut microbiome. Our study provides insightful information on the development of AD-HIES and IBD, using these mice as the basis of future fundamental, translational studies.

## Methods

### Mice

*Stat3*^fl/fl^ mice on the C57BL/6 background were originally provided by Dr. D. Levy (New York University, New York, NY) (43, 44). CD11c-Cre (*Itgax*-Cre) mice on the C57BL/6 background were purchased from The Jackson Laboratory (JAX; Bar Harbor, ME) and backcrossed to *Stat3*^fl/fl^ mice to obtain CD11c-Cre/ *Stat3*^fl/fl^ (STAT3 cKO) mice. *Rag2* KO and OT-II mice on the C57BL/6 background (both from JAX) were backcrossed to generate *Rag2* KO/OT-II mice. *IL10* KO mice on the C57BL/6 background were also obtained from JAX. The mice were bred and housed in specific pathogen-free facilities at the Indiana University School of Medicine in an AAALAC-accredited facility. Standard room environmental conditions were maintained (22+/-2°C ambient temperature, 12:12-hour light:dark cycle, and 30-70% relative humidity). Mice were fed *ad libitum* on a normal chow diet (2018SX, Envigo) or a broad-spectrum combined antibiotics-medicated diet (Uniprim, TD.06596, Envigo) where indicated. Reverse-osmosis filtered water was continuously available by auto-water lixit. Housing included autoclaved individually-ventilated cages (Max 75 cages, Alternative Design) with ¼” corncob bedding (Envigo) and tissue for enrichment. All mice were age- and sex-matched littermates; both male and female mice were used between 6 and 30 weeks of age in all experiments. All animal procedures were approved by the Indiana University School of Medicine’s Institutional Animal Care and Use Committee.

### Mouse cross-fostering

The mouse cross-fostering procedure was carried out as previously described (45, 46). *Helicobacter*-free foster breeders on a BALB/c background were purchased from JAX and housed in a separate room. Briefly, newborn litters were taken from their mother’s cage, from a known *Helicobacter*-positive room, within 24 hours of birth. After being lightly wiped with moist Rescue RTU wipes (Virox Technologies Inc.), these pups were placed into the foster mother’s cage. At the time of weaning, fecal pellets were freshly collected from these cross-fostered mice for a PCR test (IDEXX BioAnalytics, Columbia, MO) to confirm the absence of *Helicobacter spp*. in the microbiome. Additionally, mouse colony fecal samples were collected (1 fecal pellet per cage, 10 pellets pooled per test) and tested by PCR (IDEXX BioAnalytics) every 3 months as a part of the IUSM health surveillance program to confirm the *Helicobacter*-free status of the *Helicobacter*-free foster room.

### Antibodies and reagents

Allophycocyanin (APC)-, phycoerythrin (PE)– and fluorescein isothiocyanate (FITC)–conjugated monoclonal antibodies (mAbs) against murine NK cell-, DC-, B cell- or T cell-specific markers, including NK1.1, MHC class II, CD11c, B220, CD4, CD8 and TCRβ, were purchased from either BD Biosciences (San Diego, CA), or BioLegend (San Diego, CA). Fluorochrome-conjugated mAbs specific for STAT3 and IL-17 were also purchased from BD Biosciences or BioLegend. The F4/80-specific mAb was from Bio-Rad (formerly AbD Serotec). The APC-conjugated anti-FoxP3 mAb was purchased from Tonbo Biosciences (San Diego, CA). The FITC-conjugated mouse lineage marker cocktail (CD3, Gr-1, CD11b, CD45R, Ter119) was from BioLegend. APC-H7-anti-CD117 (a.k.a., c-Kit), PE-anti-CD34 and PerCP-Cy™5.5-anti-CD16/CD32 (FcγR) were purchased from BD Biosciences.

### Whole blood cell profiling

Mouse blood samples were freshly collected into EDTA-containing tubes and run through a Veterinary Hematology Analyzer (Element HT5, Heska).

### RNA/DNA extraction and qPCR

Total RNA was extracted from BMDCs using the RNeasy kit (Qiagen, Hilden, Germany) and used as a template for the synthesis of cDNA utilizing the Transcriptor First Strand cDNA Synthesis Kit (Roche, Basel, Switzerland). Total genomic DNA was extracted by boiling BMDCs in a 50 mM NaOH solution as previously described (47). Primers specific for STAT3 were designed and custom-ordered from Integrated DNA Technologies (Coralville, IA). A previously published primer pair specific for GAPDH (48) was used as an internal control. Primers for STAT3 Exons 20/21 were: Forward 5’- GAAGGAGGGGTCACTTTCACT-3’; Reverse 5’- TGCCTCCTCCTTGGGAATGTC-3’. Primers for STAT3 Introns 20/21 were: Forward 5’- CTCTCCTTCCATGGTCCGTGAC-3’; Reverse 5’- TGCCTCCTCCTTGGGAATGTC-3’. The Fast SYBR™ Green Master Mix (Thermo Fisher Scientific), together with primers, was added to each sample for the PCR reaction and run on the QuantStudio 6 Flex Real-Time PCR System (ThermoFisher Scientific). The value of STAT3 Exons 20/21 and Introns 20/21 was calculated as 2^ΔCt(GAPDH)^. All samples were analyzed in duplicate.

### Flow cytometry

Mononuclear cells were isolated from colon and liver tissues using gradient centrifugation, as previously described (49, 50). Cells from thymus, spleen, lymph nodes and Peyer’s Patches were prepared by standard procedures. Single-cell suspensions were incubated with the indicated antibodies at 4°C for 30 min. These cells were washed with HBSS containing 0.1% BSA (Sigma-Aldrich, St. Louis, MO). All cells were fixed with 1% paraformaldehyde in PBS. For intracellular cytokine staining, cells were stimulated with phorbol myristate acetate (PMA; 50 ng/ml) and ionomycin (0.5 μg/ml; Sigma-Aldrich) at 37°C; brefeldin A was added for the last 4 to 6 hours of culture. Cells were first stained with surface marker-specific antibodies, then fixed and permeabilized with 0.1% saponin and stained with cytokine-specific antibodies. To stain cells with the FoxP3-specific antibody, cells were fixed in 4% paraformaldehyde in PBS and 50% methanol in PBS, then permeabilized with 0.3% Triton X-100 in PBS. All data were acquired using an LSR4 or Fortessa cytofluorograph and analyzed using FlowJo™ v10 software (Becton Dickinson, San Jose, CA).

### Western blot analysis

STAT3 expression in different tissues was analyzed by Western blot as previously described (17). Briefly, cell lysates were resolved on an SDS-PAGE gel and transblotted onto a PVDF membrane (Millipore, Bedford, MA). The blots were probed with a STAT3-specific antibody (Cell Signaling Technology, Inc.) and then developed using chemiluminescence before exposure on film. The same blots were stripped and re-probed with antibodies against GAPDH (Cell Signaling Technology, Inc.) to ensure equal loading of proteins. Images were quantified using ImageJ (1.46v; National Institutes of Health, Bethesda, MD).

### Histology

Specimens of cecum, colon and bone (sternum or femur) were fixed with 4% neutral-buffered paraformaldehyde or 10% neutral-buffered formalin for 24-48 hours and preserved in 70% ethanol. Bones were demineralized by overnight formic acid treatment. All samples were embedded, sectioned and stained with hematoxylin and eosin (H&E) by the Indiana University School of Medicine Histology Service Core. The slides were coded and examined in a blinded fashion. A semiquantitative scoring system was used to evaluate the severity of colitis, as previously described (51).

#### Multiplex Assay

The levels of 32 different cytokines and chemokines in mouse sera were measured using a Milliplex MAP Mouse Cytokine/Chemokine premixed 32 Plex (Millipore), following the manufacturer’s protocols. The panel of magnetic beads was acquired and analyzed by a Luminex 200 instrument (Luminex Corp).

#### ELISA

To measure the levels of circulating IgE, mouse serum samples were added to rat anti-mouse IgE mAb-coated 96-well Nunc MaxiSorb plates (Thermo Scientific). An HRP-conjugated goat anti-mouse IgE antibody (SouthernBiotech) was used for detection. The levels of IL-17 were measured by using ELISA MAX^TM^ Deluxe Sets (BioLegend).

#### Microbiome analysis

Freshly-isolated mouse fecal pellets were frozen and stored in a −80°C freezer or on dry ice until analysis. Genomic DNA was isolated from fecal samples by bead-beating with the Fecal DNA Isolation Kit from Zymo Research (Irvine, CA, USA) according to the manufacturer’s instructions. The extracted DNA was immediately used for PCR or stored in standard Tris-EDTA buffer (pH 8) at 4°C. An amplicon library was constructed from isolated DNA samples by PCR to amplify the V4 region of the 16S rRNA gene with unique barcoded primers (52). Agarose gel electrophoresis was performed on the individual PCR products and visualized on a UV illuminator. The isolated PCR products were excised from the gel by the QIAquick Gel Extraction Kit (Qiagen, Germantown, MD). The NextGen sequencing Illumina MiSeq platform was used for sequencing the PCR products of about 250 bp paired-end reads from the V4 region of the 16S rRNA gene (52).

The obtained raw FASTQ files were used for library construction, de-multiplexed, and assessed for quality control using FastQC. An automated pipeline, QWRAP, was developed that runs all of these tools (52). This pipeline has recently been extended using DADA2 to provide a more robust error model supporting sample filtering and clustering (53). Further analysis of the beta diversity metrics (weighted UniFrac, unweighted Unifrac and Bray Curtis) were determined as previously described (54).

#### Statistical analysis

A Kruskal–Wallis test was performed to identify key microbiome taxa with changes in relative abundances between groups. These statistical tests were performed using tools within the QIIME package. Graphs were generated and statistical analyses were performed using GraphPad Prism 8. Groups were compared using the Mann–Whitney unpaired *t* test or one-way ANOVA. The data are shown as the mean ± standard deviation. The log-rank test was performed for survival analysis. *p* values <0.05 were considered statistically significant.

## Author contributions

JL and RRB conceived the project. AKI, MO, JD, BZ, MHK and ALD devised the methodology, JL, KLS, MC, WVDP, CM and JH performed the experiments and analyzed the data along with RRB. JL, RRB, KLS, MC, CM wrote the manuscript.

## Acknowledgements

The authors would like to thank the Indiana University Simon Cancer Center Flow Cytometry Facility, Histology Service Core at the Indiana University School of Medicine (IUSM), and Hong Liu for technical assistance, as well as Drs. George Sandusky, Xinxin Huang and Dongyeop Shin for helpful discussions. This work was supported by a Showalter Research Trust Fund award (to JL), funds from the Department of Microbiology & Immunology at IUSM (to RRB), U54 DK106846 from the NIH supporting IUSM’s Cooperative Center of Excellence in Hematology, and UL1TR001417 from the NIH National Center for Advancing Translational Sciences (NCATS) supporting the UAB Center for Clinical and Translational Science.

## Supplemental Figure Legends

Fig. S1: STAT3 expression is deficient in BMDCs from STAT3 cKO mice. A) An illustration of the genomic structure of floxed sequences of the *stat3* gene in STAT3 cKO mice. Solid triangles indicate the loxP sites. Short arrows indicate the qPCR primers for exon 20, intron 20/21 and exon 21 of the *stat3* gene. B) Total RNA was extracted from BMDCs and cDNA was generated. The expression level of exon 20/21 was measured by qPCR. C) Genomic DNA was extracted from BMDCs and the level of expression of introns 20/21 and exons 20/21 was measured by qPCR. D) B220-positive spleen cells were sorted using anti-B220 mAb-conjugated magnetic beads. Expression of STAT3 in enriched B cells (E) and thymocytes (F) was measured by Western blot analysis. Each dot represents an individual animal. *, *p*<0.05; Mann–Whitney unpaired *t* test.

Fig. S2: Altered leukocyte subpopulation levels in STAT3 cKO mice. (A) Gating strategy for DCs, macrophages, NK, B and T cells. (B) Splenic lymphocytes (T, NK and B cells) are reduced in STAT3 cKO mice. (C) DCs and macrophages from the spleen are present at higher numbers in STAT3 cKO than in WT/het mice. (D) Gating strategy for identifying Th1 and Th17 cells. (E) STAT3 cKO mice have similar levels of Th1 cells as WT mice. (F) STAT3 cKO mice have comparable levels of Th17 cells in the Peyer’s Patch as WT mice. (G) Gating strategy for identifying Treg cells. (H) STAT3 cKO mice have WT levels of Treg cells. *, *p*<0.05; **, *p*<0.01; ***, *p*<0.001. Mann–Whitney unpaired *t* test.

Fig. S3: Altered gut microbiome in GM-IUSM STAT3 cKO mice. Fecal pellets from GM-IUSM and GM-JAX STAT3 cKO mice and their WT/het littermates were analyzed by 16S rRNA sequencing analysis. The Amplicon Sequence Variants (ASV) of *Pseudomonadaceae* and *Anaeroplasmataceae* (A), *Deferribacteraceae*, *Helicobacteraceae* and *Mycoplasmataceae* (B), *Rikenellaceae* and *Prevotellaceae* (C) and, *Enterococcaceae* and *Enterobacteriaceae* (D) are shown. The data are plotted as the mean ± SD. Each dot represents an individual animal. *, *p*<0.05; ***, *p*<0.001. One-way ANOVA test.

Fig. S4: IL-10 KO mice do not have elevated IL-17 or IgE in the circulation. (A and B) Serum IgE and IL-17 from WT/het, STAT3 cKO and IL-10 KO mice are shown. Each dot represents an individual animal. NS, not significant; *, *p*<0.05; **, *p*<0.01; ***, *p*<0.001. One-way ANOVA test.

