## Supplemental Figures for "STAT3 expression in dendritic cells protects mice from colitis by a gut microbiome-dependent mechanism"

Liu et al
Fig. S1

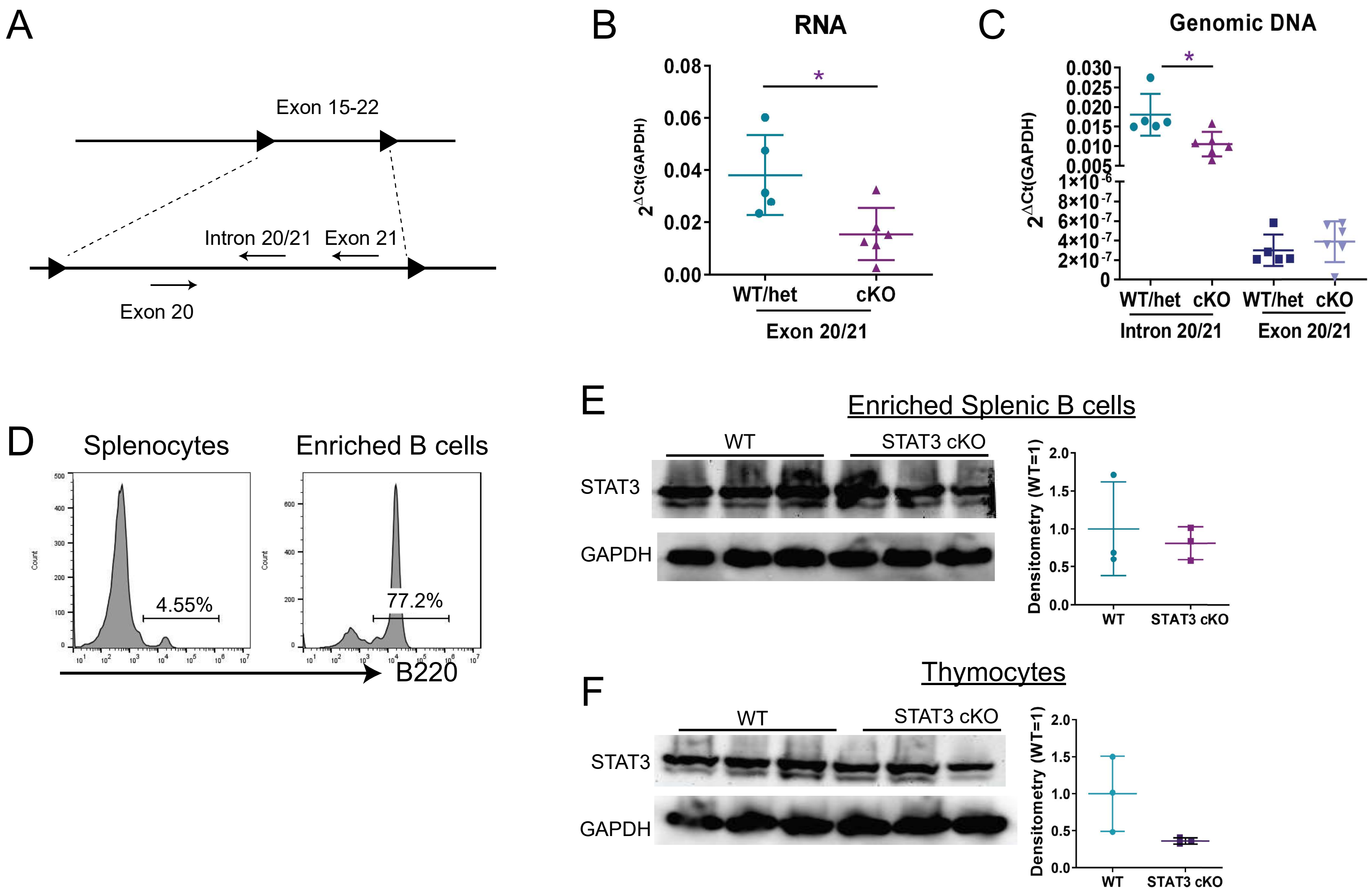


Supplemental Figure 1. STAT3 expression is deficient in BMDCs from STAT3 cKO mice. (A) An illustration of the genomic structure of floxed sequences in the *Stat3* gene of STAT3 cKO mice. Solid triangles indicate loxP sites. Short arrows indicate the qPCR primers for exon 20, intron 20/21 and exon 21 of the Stat3 gene. (B) Total RNA was extracted from BMDCs and cDNA was generated. The expression level of exon 20/21 was measured by qPCR. (C) Genomic DNA was extracted from BMDCs and the level of expression of introns 20/21 and exons 20/21 was measured by qPCR. (D) B220-positive spleen cells were sorted using anti-B220 mAb-conjugated magnetic beads. Expression of STAT3 in enriched B cells (E) and thymocytes (F) were measured by Western blot analysis. Each dot represents an individual animal. *, $p$<0.05, Mann–Whitney unpaired $t$ test.

Liu et al
Fig. S2

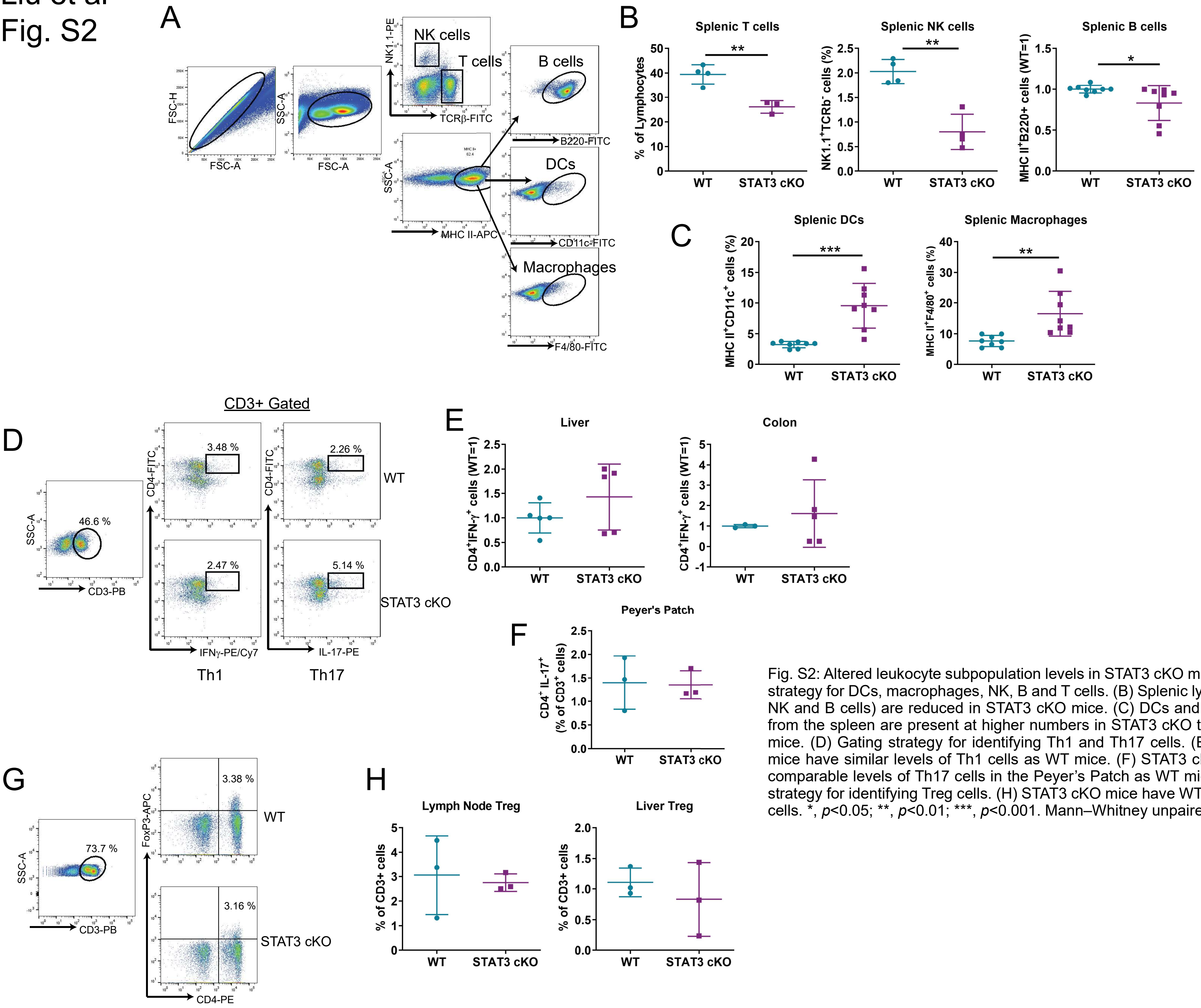


Fig. S2: Altered leukocyte subpopulation levels in STAT3 cKO mice. (A) Gating strategy for DCs, macrophages, NK, B and T cells. (B) Splenic lymphocytes (T, NK and B cells) are reduced in STAT3 cKO mice. (C) DCs and macrophages from the spleen are present at higher numbers in STAT3 cKO than in WT/het mice. (D) Gating strategy for identifying Th1 and Th17 cells. (E) STAT3 cKO mice have similar levels of Th1 cells as WT mice. (F) STAT3 cKO mice have comparable levels of Th17 cells in the Peyer's Patch as WT mice. (G) Gating strategy for identifying Treg cells. (H) STAT3 cKO mice have WT levels of Treg cells. *, $p$<0.05; **, $p$<0.01; ***, $p$<0.001. Mann–Whitney unpaired $t$ test.

Liu et al
Fig. S3

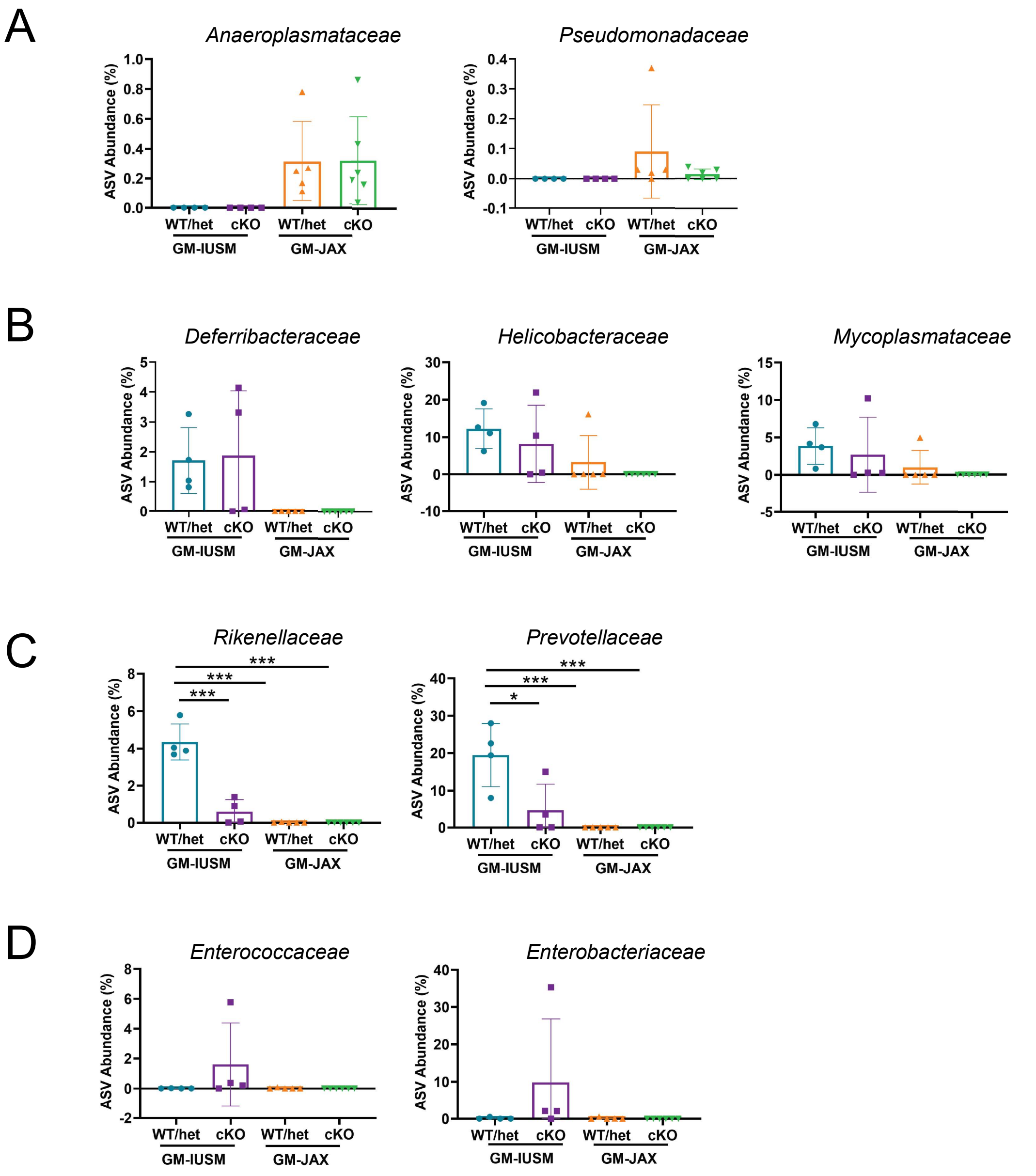


Fig. S3: Altered gut microbiome in GM-IUSM STAT3 cKO mice. Fecal pellets from GM-IUSM and GM-JAX STAT3 cKO mice and their WT/het littermates were analyzed by 16S rRNA sequencing analysis. The Amplicon Sequence Variants (ASV) of *Pseudomonadaceae* and *Anaeroplasmataceae* (A), *Deferribacteraceae*, *Helicobacteraceae* and *Mycoplasmataceae* (B), *Rikenellaceae* and *Prevotellaceae* (C) and, *Enterococcaceae* and *Enterobacteriaceae* (D) are shown. The data are plotted as the mean ± SD. Each dot represents an individual animal. *, $p<0.05$; ***, $p<0.001$. One-way ANOVA test.

Liu et al
Fig. S4

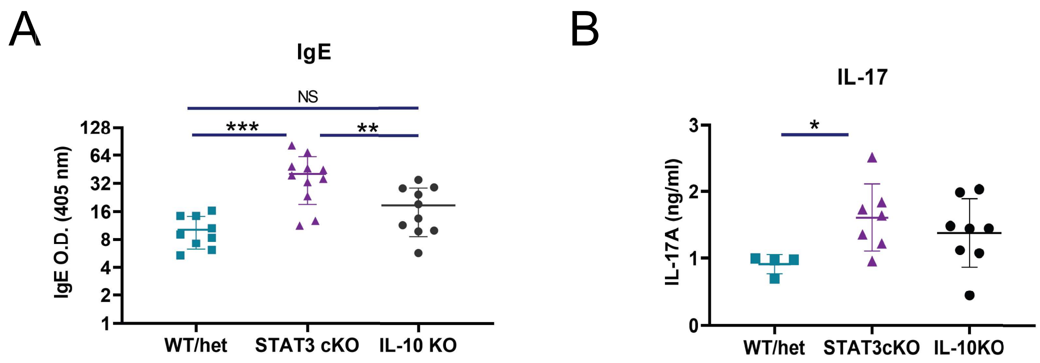


Fig. S4: IL-10 KO mice do not have elevated IL-17 or IgE in the circulation. (A and B) Serum IgE and IL-17 from WT/het, STAT3 cKO and IL-10 KO mice are shown. Each dot represents an individual animal. NS, not significant; *, $p$<0.05; **, $p$<0.01; ***, $p$<0.001. One-way ANOVA test.
